# Ribosomal RNA methylation by GidB modulates discrimination of mischarged tRNA

**DOI:** 10.1101/2021.03.02.433644

**Authors:** Zhuo Bi, Yu-Xiang Chen, Iris D. Young, Mohamad T. Dandan, Hemant Joshi, Hong-Wei Su, Yuemeng Chen, Jia-Yao Hong, James S. Fraser, Babak Javid

**Author notes:** These authors contributed equally (Zhuo Bi, Yu-Xiang Chen, Iris D. Young).

## Abstract

Despite redundant cellular pathways to minimize translational errors, errors in protein synthesis are common. Pathways and mechanisms to minimize errors are classified as pre-ribosomal or ribosomal. Pre-ribosomal pathways are primarily concerned with the appropriate charging of tRNAs with their cognate amino acid. By contrast, the ribosomal decoding centre is considered ‘blind’ to mischarged tRNAs since these have cognate codon•anti-codon pairing. Here, we identified that in mycobacteria, deletion of the 16S ribosomal RNA methyltransferase *gidB* led to increased ribosomal discrimination of mischarged tRNAs. Discrimination only occurred in mycobacteria enriched from environments or genetic backgrounds with high rates of mistranslation. GidB deletion was necessary but not sufficient for reducing mistranslation due to misacylation. Analysis of new cryoEM structures of the *M. smegmatis* ribosomes derived from wild-type and *gidB*-deleted strains point to the interaction between the base methylated by GidB on the 16S RNA and an asparagine on the ribosomal S12 protein that when mistranslated to aspartate may be involved in altering translational fidelity. Our data suggest a mechanism by which mycobacterial ribosomes can discriminate mischarged tRNAs and that 16S rRNA differential methylation by GidB may act to prevent catastrophic translational error.

## Introduction

All clades of life have evolved multiple, redundant pathways to reduce translational error [1, 2]. Despite these mechanisms, errors in protein synthesis are remarkably common and are orders of magnitude more frequent than errors in DNA or RNA synthesis [3–7]. There is no one ‘optimal’ rate of error. Both the error rates and the dominant sources of error can be both species and even organelle specific [1, 8]. Furthermore, translational errors (mistranslation) may result in adaptive phenotypes, particularly in the context of environmental stressors [1, 5, 6, 9–20]. However, excess mistranslation can also cause protein aggregation [21, 22], organ degeneration [23, 24] and is the mechanism for the bactericidal activity of aminoglycosides [25]. Collectively, these lines of evidence suggest that no optimal balance for translational error exists and that selection favors tunable and context-specific mistranslation rates [1, 3, 5, 26]. However, the precise mechanisms by which mistranslation rates are tuned are poorly understood.

In addition, molecular mechanisms of translational error vary considerably, as do the proof-reading pathways that have evolved to reduce them. Generally, sources of error and proof-reading can be divided into pre-ribosomal and ribosomal mechanisms [1]. Pre-ribosomal errors arise from mischarging of the tRNA with a non-cognate amino acid and proof-reading mechanisms include pre-and post-transfer editing functions of aminoacyl tRNA synthetases [27]. Following aminoacyl-tRNA synthesis, *trans*-acting editing mechanisms can reject mischarged tRNAs [28, 29], and the aminoacyl-tRNA chaperone, EF-Tu optimally binds cognate aminoacyl-tRNAs compared with mischarged tRNAs [30]. These multiple and redundant pre-ribosomal proof-reading steps – solely concerned with the charging of tRNA with its cognate amino acid – have been proposed as necessary since previously described ribosomal proof-reading mechanisms ensure cognate codon·anticodon pairing [2, 31] and therefore are insensitive to the identity of the amino acid charged to the tRNA.

In mycobacteria, the dominant source of translational error is due to the pre-ribosomal indirect tRNA aminoacylation pathway [12, 32, 33]. Most bacteria, with the notable exception of a few proteo-bacteria such as *Escherichia coli,* lack either the glutaminyl- or asparaginyl-tRNA synthetases, or both [34]. Instead, a two-step indirect tRNA synthesis pathway is required to ensure cognate charging of glutamine and asparagine tRNAs. In the first step, a non-discriminatory glutamyl or aspartyl synthetases mischarge glutaminyl-tRNA with glutamate (i.e. Glu-tRNA^Gln^) and asparaginyl-tRNA with aspartate (i.e. Asp-tRNA^Asn^) respectively. In the second step, the enzyme GatCAB corrects the mischarged aminoacyl moiety by transamidation [35] and see Fig. 1A. Despite its essential function, partial loss-of-function mutations in *gatCAB* can be readily selected in mycobacteria, which lacks both synthetases and mutations in *gatCAB* occur naturally in clinical isolates of pathogenic *M. tuberculosis* [12, 36, 37]. These mutant strains exhibit extremely high (up to 10%/codon) rates of the mistranslation of Gln/Asn codons to Glu/Asp, respectively and specifically [12]. Even in wild-type mycobacteria, the specific mistranslation rates of Gln→Glu and Asn→Asp, measured using similar gain-of-function reporters, are orders of magnitude higher than in *E. coli* – that has the full set of 20 aminoacyl tRNA synthetases and hence lacks GatCAB [38].

**Figure 1.**
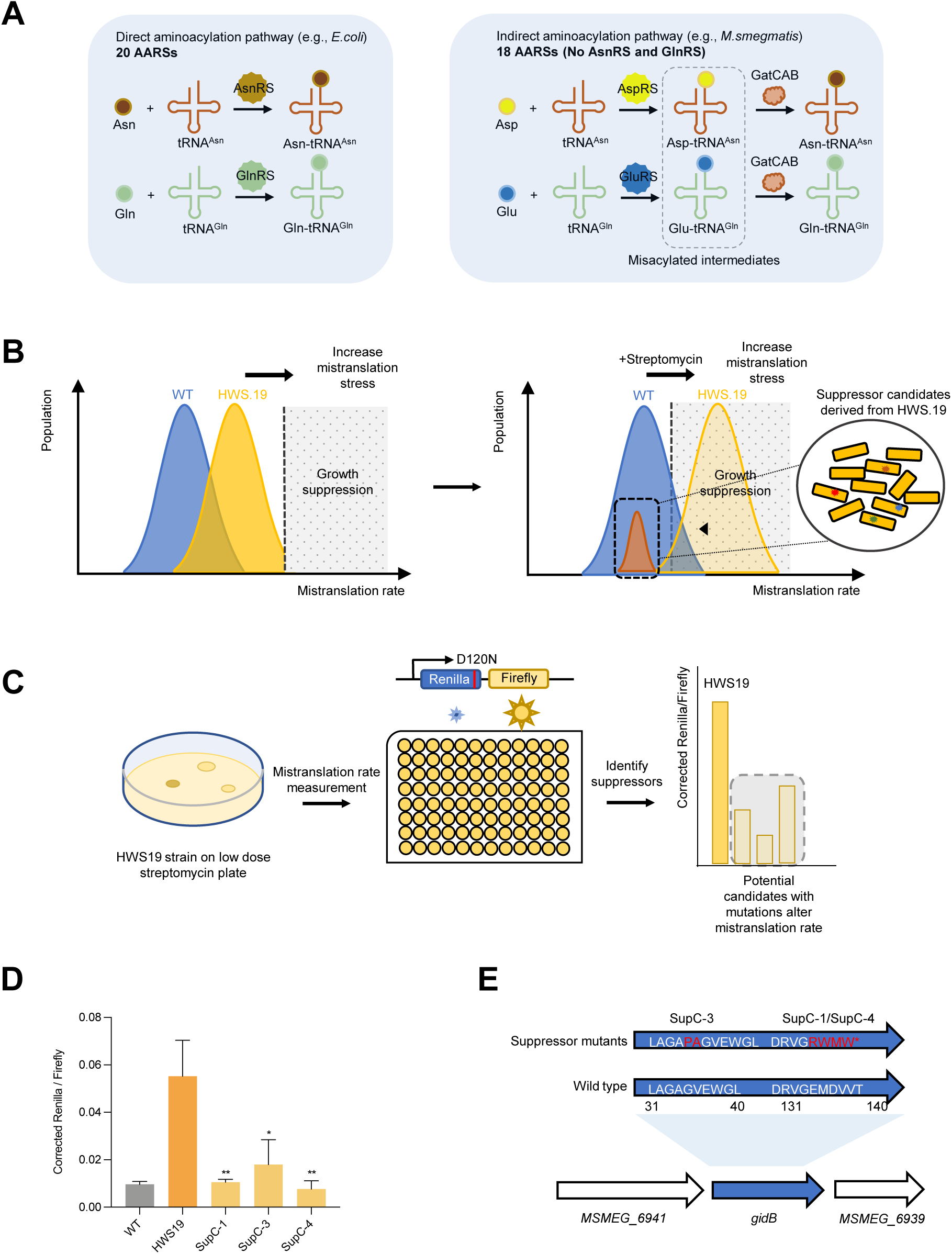
Suppressor screen identifies gidB as a potential fidelity factor. **A.** Indirect aminoacylation pathway for tRNA^Asn^ and tRNA^Gln^ in mycobacteria. **B**. Design of the suppressor screen: applying an increasing mistranslation stress to both strains makes HWS.19 more susceptible to stress and more likely to have mutation in suppressors. **C.** Workflow of suppressor screen. **D**. N to D mistranslation rate of suppressor candidates compared with WT and HWS.19 measured by Renilla-Firefly dual luciferase reporter as depicted in Figure1C. **E.** *gidB* mutation was mapped onto 3 candidates by whole genome sequencing. (*P < 0.05, **P < 0.01, ***P < 0.001, Student *t* test)

Here, we investigated whether further fidelity factors can be identified in mycobacteria. We identified that deletion of the 16S ribosomal RNA (rRNA) methyltransferase *gidB* is necessary, but not sufficient for the discrimination of mischarged tRNAs. Deletion of *gidB* only increased discrimination of mischarged tRNAs in mycobacteria with elevated mistranslation rates – due to mutation or environmental context. Solving the structure of mycobacterial ribosomes purified from +/-*gidB* strains suggested that methylation of rRNA may interact with mistranslated ribosomal amino acids in the small subunit, implying a potential mechanism by which rRNA methylation contributes to fidelity. Collectively, these results point to an active ribosomal proof-reading of ribosomes beyond codon•anticodon pairing and suggest that nonmethylation of rRNA prevents catastrophic translational error.

## Results

### A suppressor screen in mycobacteria identifies *gidB* as a potential translational fidelity factor

We previously used forward genetic screens to identify strains of *Mycobacterium smegmatis* with extremely high specific rates of mistranslation – i.e. mistranslation of Asn→Asp. These screens identified mutations in the essential amidotransferase genes *gatCAB* that resulted in high rates of mistranslation of Gln/Asn, specifically [12, 36]. We hypothesized that the mycobacterial genome may encode for other ‘fidelity factors’ that could modulate mistranslation rates in the background of a compromised indirect tRNA aminoacylation pathway. We, therefore, designed a suppressor screen strategy in the strain HWS19 [12], which encodes a three amino acid deletion in *gatA* and, consequently, has extremely high rates of translational error of Gln→Glu and Asn→Asp specifically (Fig 1B). We wondered whether HWS19, with a high background mistranslation rate of Gln/Asn codons, is more susceptible to aminoglycosides, such as streptomycin that are known to increase ribosomal decoding errors across multiple codons [33, 39]. Plating of equivalent colony forming units (CFU) of strain HWS19 on low-dose streptomycin-agar led to the recovery of significantly fewer colonies compared with wild-type (Fig. S1), confirming HWS19 was hypersusceptible to streptomycin. The basis of our suppressor screen, therefore, was as follows: plating of HWS19 onto low-dose streptomycin-agar should select for strains with mutations that *decrease* mistranslation on this high mistranslating background (Fig. 1B, 1C). Suppressors identified from selection on low-dose streptomycin agar were then tested using our custom dual-luciferase mistranslation reporter system to verify that they had decreased error rates (Fig. 1C).

To identify mutants leading to decreased mistranslation, we plated 1×10^7^ CFU onto each of 6 agar plates containing 1 µg/mL streptomycin. The remaining survivors were transformed with dual-luciferase reporter plasmids that measured specific mistranslation errors of asparagine for aspartate [12, 17, 40] to identify low mistranslating candidates (Fig. 1C). We initially identified 4 suppressor mutants with lower mistranslation rates compared with the parent HWS19 strain, and comparable to wild-type *M. smegmatis* (Fig. 1D). Sequencing the strains revealed three of the four had mutations in the 16S rRNA methyltransferase *gidB* (Msmeg_6940) – Table S1. The remaining suppressor (C-23) had multiple mutations including in the gene, *tuf*, coding for the aminoacyl-tRNA chaperone EF-Tu. (Table S1). We subsequently sequenced just the *gidB* gene of a further 10 candidate suppressors and identified a further seven with a total of three additional independent mutations in *gidB* (Table S2). All but one of these further suppressors also had reduced mistranslation rates (Fig. S2), strongly supporting that mutations in *gidB* are able to suppress the high specific mistranslation rates in strain HWS19 that arise from mischarging tRNA.

### Deletion of *gidB* increases ribosomal discrimination against misacylated tRNA in mycobacteria with high mistranslation

Loss of function mutations in GidB can cause low-level streptomycin resistance: lack of methylation in 16S rRNA interferes with streptomycin binding [41, 42]. We were, therefore, concerned that our candidates from the screen may simply represent selection against streptomycin. To determine whether the observed phenotype was independent of streptomycin selection, we deleted *gidB* in both the HWS19 and parental (wild-type) *M. smegmatis* backgrounds in the complete absence of streptomycin selection. Deletion strains were complemented with a chromosomally integrated plasmid expressing *gidB* from its native promoter (Fig. S3). We measured specific mistranslation rates using two complementary reporter systems: the dual-luciferase system, used in the initial screen, as well as a dual-fluorescent reporter system (Fig. S4, S5) that uses flow cytometry to measure relative mistranslation rates [12]. This reporter, a GFP-mRFP fusion protein had a glutamate necessary for GFP fluorescence mutated to glutamine, abrogating green, but not red fluorescence [12]. Increased specific mistranslation of Gln→Glu would result in increased GFP fluorescence and hence increased GFP/RFP ratios. Measurement of specific mistranslation rates with both reporters verified that deletion of *gidB* significantly increased translational fidelity in strain HWS19, and this phenotype could be readily complemented (Fig. 2A, B, left panel). Surprisingly, deletion of *gidB* had no phenotype using the same reporters on a wild-type background (Fig. 2A, B, right panel).

**Figure 2.**
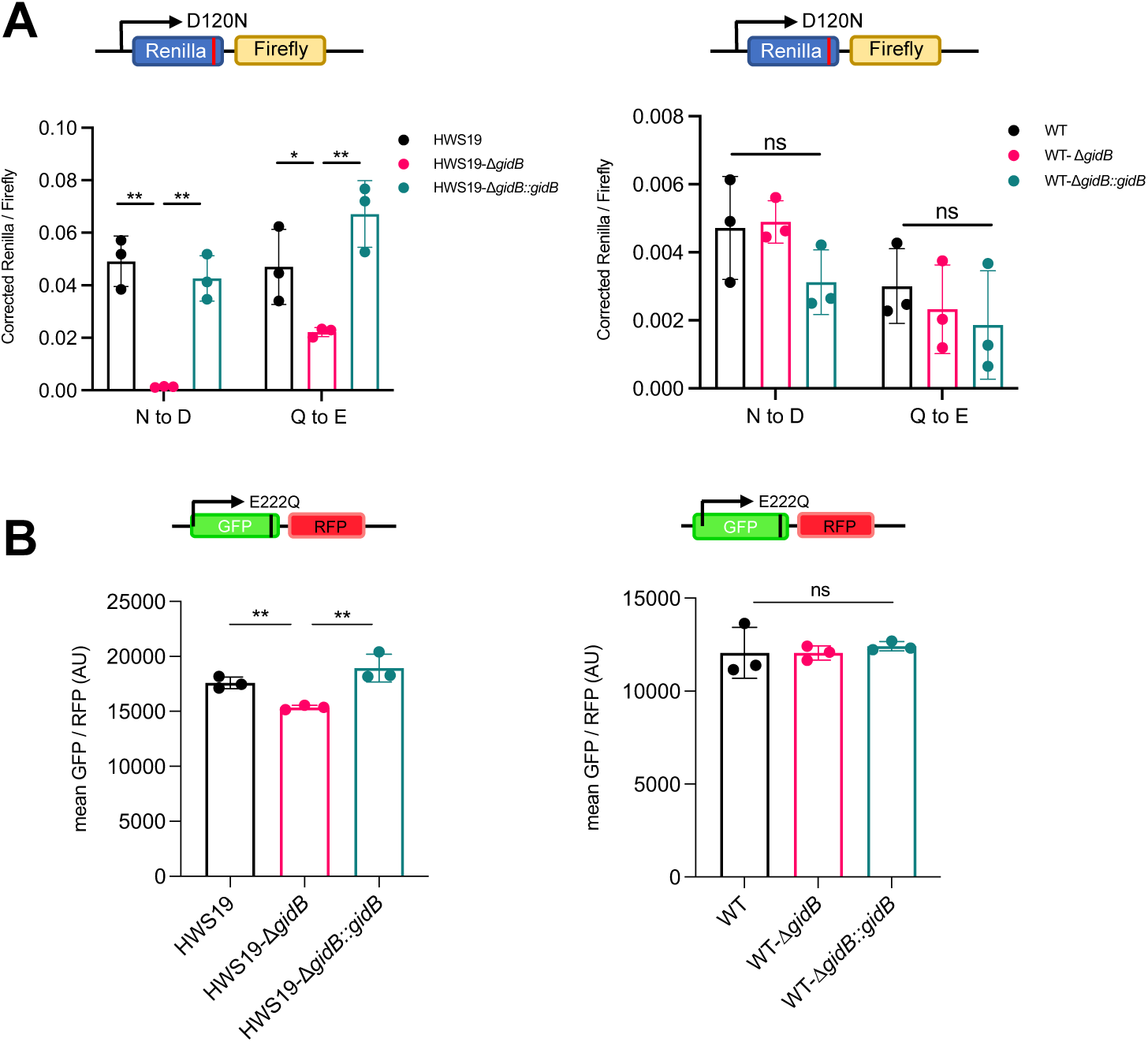
Deletion of GidB in mycobacteria increases translation fidelity in high mistranslation mutant. **A.** N to D Mistranslation rate measured by gain-of-function Renilla-Firefly dual luciferase reporter in high mistranslation HWS19 strain (Left) and wild-type strain (Right). **B.** Q to E Mistranslation rate measured by gain-of-function dual fluorescent reporter in high mistranslation HWS19 strain (Left) and wild-type strain (Right).

### GidB is necessary for discrimination of mischarged tRNA under conditions that enrich for relatively high mistranslation rates

Our results suggested that deletion of *gidB* increased translational fidelity only in a mutant strain (HWS19) that had basal extremely high rates of mistranslation arising from lack of quality control of mischarged asparagine and glutamine tRNAs and that deletion of *gidB* had no phenotype in wild-type mycobacteria grown under standard laboratory conditions. We wondered whether deletion of *gidB* had a phenotype in wild-type mycobacteria, but only under conditions associated with increased rates of mistranslation. We had previously shown that exposure of wild-type mycobacteria to rifampicin at 1x minimal bactericidal concentration (MBC) could select for survival and growth of *wild-type* mycobacteria due to two reversible and non-genetic mechanisms: increased rates of specific mistranslation [12] and a semi-heritable survival program [43], and not due to mutations in the rifampicin’s target *rpoB* or other mutations. We plated wild-type *M. smegmatis* on non-selective LB medium or low-dose (1xMBC) rifampicin-agar (**Fig. 3A**). As before, deletion of *gidB* isolated from non-selective medium had no phenotype (**Fig. 3B**). However, bacteria that survived and grew on rifampicin-agar had increased rates of mistranslation as measured by the dual-fluorescent reporter, in keeping with prior observations [12, 43]. These increased mistranslation rates were reverted by deletion of *gidB* in fully complementable manner (**Fig. 3B**), suggesting that deletion of *gidB* increased translational fidelity under growth conditions causing high rates of mistranslation, even in wild-type bacteria.

**Figure 3.**
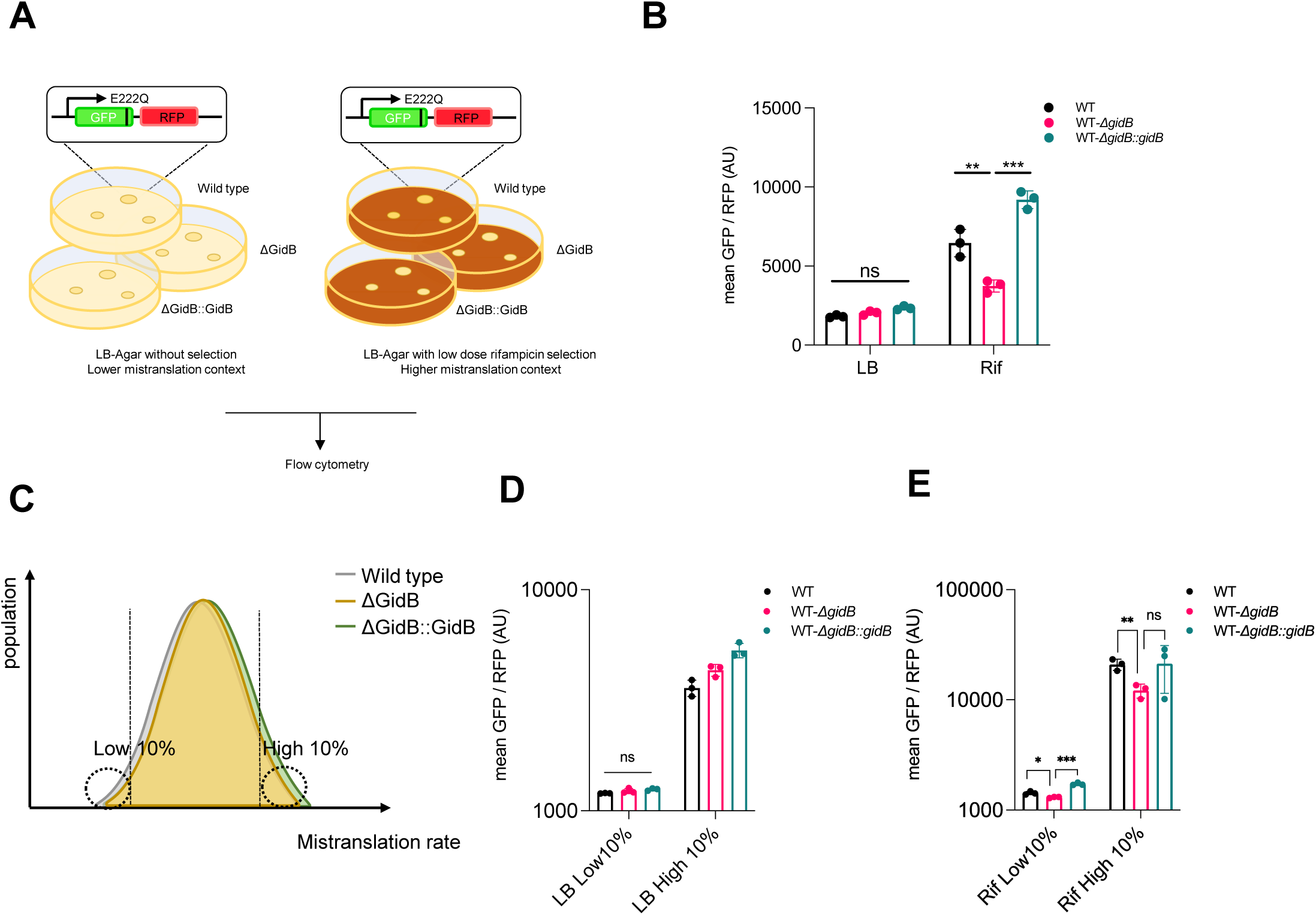
Deletion of GidB in mycobacteria increases translation fidelity under high mistranslation physiological context. **A.** Schematic of measuring mistranslation rate using gain-of-function dual fluorescence reporter under normal and high mistranslation physiological context **B.** Q to E mistranslation of mycobacteria scraped from LB-agar or LB-agar with low dose rifampicin. **C.** Gating strategy to gate top/bottom 10% of bacteria with high/low mistranslation rate. **D.** Q to E mistranslation of different bacteria population (no selection) gated from the highest/lowest mistranslation rate as described in C. **E**. Q to E mistranslation of different bacteria population (Rif selection) gated from the highest/lowest mistranslation rate as described in C (*P < 0.05, **P < 0.01, ***P < 0.001, Student *t* test)

The use of a dual-fluorescent reporter and flow cytometry for measurement of relative mistranslation rates allowed us to measure the impact of *gidB* deletion on different subpopulations of mycobacteria in the same experiment: i.e. we could selectively gate on heterogeneous subpopulations that had relatively high or low GFP/mRFP ratios, representing high and low mistranslation rates specifically (**Fig. 3C**). This allowed us to ask the question as to whether deletion of *gidB* was sufficient, as well as necessary, for the discrimination of mischarged tRNA. We gated on bacteria with the highest (10%) and lowest (10%) green/red (i.e. relative mistranslation rates) ratios from the two experimental conditions, i.e., bacteria isolated from ‘standard’ non-selective LB agar, and from low-dose rifampicin agar, which enriched for bacteria with higher average mistranslation rates. Deletion of *gidB* had no observable phenotype on translational fidelity in bacteria isolated from non-selective media, even when only the cells with the highest apparent mistranslation rates were analysed (**Fig. 3D**). By contrast, in bacteria enriched for higher mistranslation rates (rifampicin exposure) *gidB* deletion could revert the mistranslation phenotype in a complementable manner (**Fig. 3E**). Therefore, *gidB* deletion appears necessary, but not sufficient for translational fidelity, and it is only within environmental contexts that enrich for high rates of mistranslation that deletion of *gidB* causes increased ribosomal discrimination of misacylated tRNA in otherwise wild-type mycobacteria.

For ribosomal decoding, there is consensus that there is a trade-off between speed versus accuracy [44, 45]. We first measured growth rates of both WT *M. smegmatis* and HWS19 strains with/without deletion of *gidB* to determine if the deletion impacted general fitness. Growth rates in both cases were identical between the “wild-type”, *gidB* deletion and complemented strains (**Fig. S6**). We then measured translational efficiency through induction of Nluc luciferase and measurement of Nluc activity, as before [32]. Deletion of *gidB* had no impact on efficiency of Nluc translation in either strain, suggesting that in axenic culture, lack of GidB does not impact rate of synthesis of this protein (**Fig. S7**).

### Testing the generality of mischarged tRNA discrimination by deletion of *gidB* beyond mischarged tRNA^Gln^ and tRNA^Asn^

The previous experiments confirm that deletion of *gidB* increased discrimination of physiologically mischarged Glu-tRNA^Gln^ and Asp-tRNA^Asn^. We next wanted to ask whether this discrimination of mischarged tRNA was a general property that would allow discrimination of any mischarged tRNA. To test this idea, we leveraged the unique properties of tRNA^Ala^ and their cognate synthetase. To ensure the accurate charging of tRNAs, most aminoacyl tRNA synthetases depend on recognizing the anticodon of their cognate tRNAs as an identity element. For alanyl synthetase (AlaRS), the sole identity element in all clades of life is a G^3^•U base-pair in the tRNA acceptor stem, and AlaRS can charge any tRNA with this identity element, regardless of anti-codon identity, i.e. the G^3^•U base-pair is both necessary and sufficient for alanine aminoacylation by AlaRS [46–48]. Therefore, the anticodon of any alanyl-tRNA can be mutated to any triplet, and the tRNA will still be charged with aminoacyl-alanine [46] and will mediate specific translational errors. We had previously mutated the anticodon of a mycobacterial alanyl-tRNA to CCA, coding for tryptophan, which would result in specific mistranslation of alanine for tryptophan at UGG codons [17] – **Fig. 4A**. To measure specific mistranslation, we modified the dual-luciferase reporter system. We identified a critical alanine residue in Renilla luciferase, which, when mutated to tryptophan (A214W), resulted in >20-fold decrease in activity (**Fig. S8**). This reporter would thus be able to discriminate mistranslation errors greater than 5%/codon of tryptophan to alanine. We transformed the mutant alanyl-tRNA (tRNA_CCA_^Ala^), cloned into a tetracycline-inducible plasmid into wild-type and Δ*gidB M. smegmatis*, along with the specific dual-luciferase reporters. Induction of tRNA_CCA_^Ala^ resulted in increased specific mistranslation of tryptophan for alanine, as expected. Deletion of *gidB* did not increase translational fidelity, even in this high mistranslating context (**Fig. 4B**). Next we transformed the mutant alanyl-tRNA and reporters into The high basal mistranslation (Gln→Glu, Asn→Asp, specifically) strain HWS19 and measured the rates of Trp→Ala mistranslation using the new Ren-FF dual-luciferase reporter. Deletion of *gidB* decreased rates of Trp→Ala mistranslation as measured by rescue of Renilla activity in the HWS19 background, suggesting that in this context, Δ*gidB* ribosomes could discriminate against even Trp→Ala mistranslation (**Fig. 4C**). Surprisingly, this phenotype is only observed in this strain with high rates of ‘physiological’ mistranslation and not in the WT strain. Taken together, these results suggested deletion of *gidB* was necessary but not sufficient for discrimination of mischarged tRNAs and that a further environmental context, possibly generated via excess ‘physiological’ mistranslation or some other stressor was also required.

**Figure 4.**
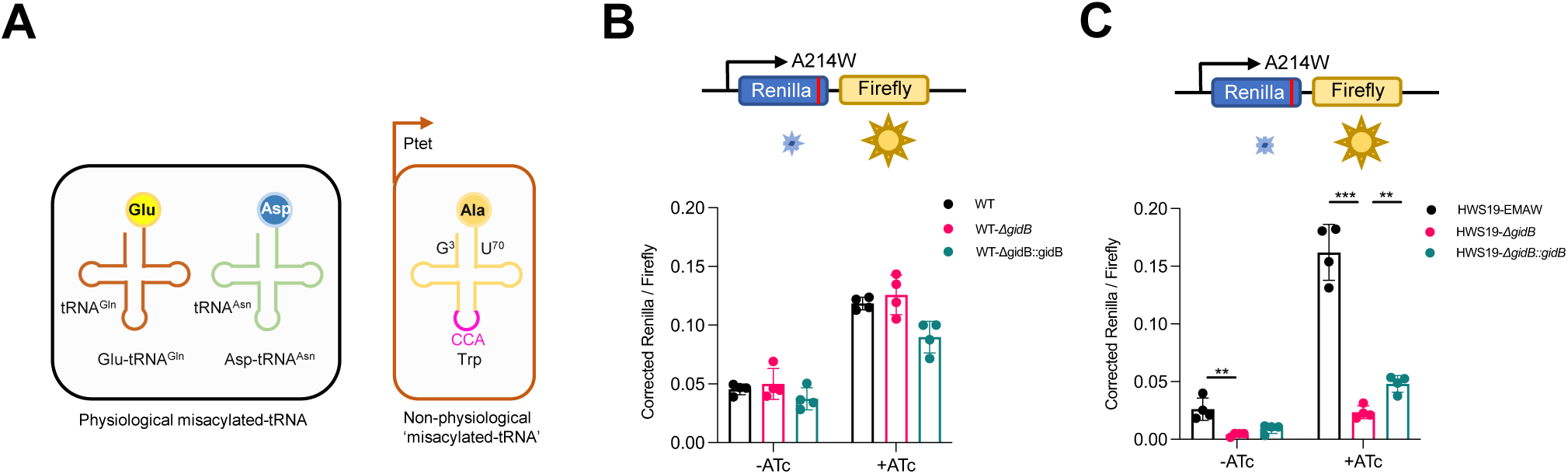
Deletion of GidB decreases mistranslation mediated by non-physiological ‘misacylated-tRNA’. **A.** The anticodon of an alanine-tRNA is mutated to tryptophan codon (EMAW) result in mistranslation from tryptophan to alanine (Right) to mimic non-physiological miacylated-tRNA compared to the two natural misacylated-tRNAs (Left) in mycobacteria. Mistranslation rate of tryptophan to alanine measured by Renilla-Firefly reporter (top) in high mistranslation HWS19 strain (**B**) and wild-type strain (**C**) with or without EMAW expression. (*P < 0.05, **P < 0.01, ***P < 0.001, Student *t* test)

### Deletion of *gidB* causes reduced tolerance to rifampicin

We tested the physiological relevance of increased translational fidelity caused by *gidB* deletion in rifampicin antibiotic tolerance. We had previously demonstrated that increased, specific, mistranslation due to Glu-tRNA^Gln^ and Asp-tRNA^Asn^ mischarging caused increased tolerance to the antibiotic rifampicin due to mistranslation of critical residues in the drug target, RpoB [12, 17, 36]. Since antibiotic tolerance is mediated by a subpopulation of bacteria that resist killing [49–59], we hypothesised that deletion of *gidB*, and the subsequent increase in translational fidelity, would render the most tolerant subpopulation susceptible to rifampicin. Deletion of *gidB* resulted in significantly increased killing of *M. smegmatis* exposed to rifampicin in axenic culture in both the high mistranslating strain HWS19 and, crucially, also in wild-type mycobacteria (**Fig. 5A, B**). This suggests that rifampicin treatment is a sufficient condition for deletion of *gidB* to exert its effect.

**Figure 5.**
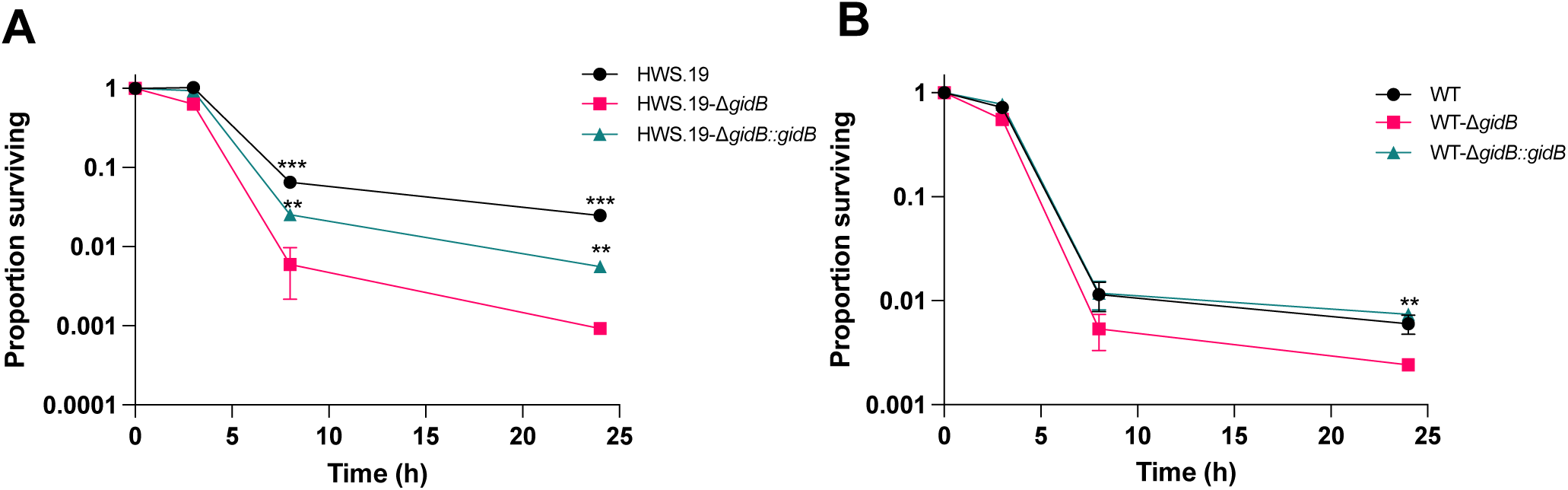
Deletion of GidB decreases rifampicin tolerance. Rifampicin killing curve in high mistranslation background (A) and wild-type background (B). T.test was performed between *gidB* deletion strain and the other two strains respectively. (*P < 0.05, **P < 0.01, ***P < 0.001, Student *t* test)

### Molecular mechanism of GidB-mediated methylation in the ribosome

As a first step to identify the molecular mechanism of changes in adaptive mistranslation, we wished to resolve the structure of ribosomes from HWS19 and HWS19-Δ*gidB* cells for analysis by cryo-electron microscopy. Sucrose gradient ultracentrifugation and ribosome subunit profiling of ribosomes from the strains did not reveal difference in the distribution of different ribosome species, suggesting that deletion of *gidB* did not alter ribosome biogenesis (**Fig. S9**). Three dimensional reconstructions of 70S fractions isolated from sucrose gradient and the models refined into those maps did not reveal significant changes in structure, likely due to the limited resolution (**Fig. 6**). Analysis of the resulting models does point to a hypothesis for the modulation of mistranslation in Δ*gidB* ribosomes as a function of GatCAB activity: an interaction between 16S G507 (the target of GidB methylation) and S12 Asn46 (**Fig. 6**). When GatCAB functions, minimizing mistranslation, as in WT *M. smegmatis*, Asn46 is reliably coded as Asn; however, in the HSW19 background and other conditions with diminished GatCAB activity, Asn46 will be coded as Asp at a much higher frequency. The interaction between G507 and the S12 loop containing Asn46 has previously been implicated in stabilizing tRNA-mRNA interactions during aminoacyl-tRNA selection in the decoding process in *M. tuberculosis* [42]. Phenotypic misincorporation from Asn to Asp likely alters this interaction and the conformational dynamics of this key loop. Further structural work, at higher resolution and including structures of translating ribosomes, will be needed to unravel how the interactions between the G507 methylation, tRNA identity, mRNA structure, and antibiotic binding affect the relative rates of adaptive mistranslation.

**Fig. 6.**
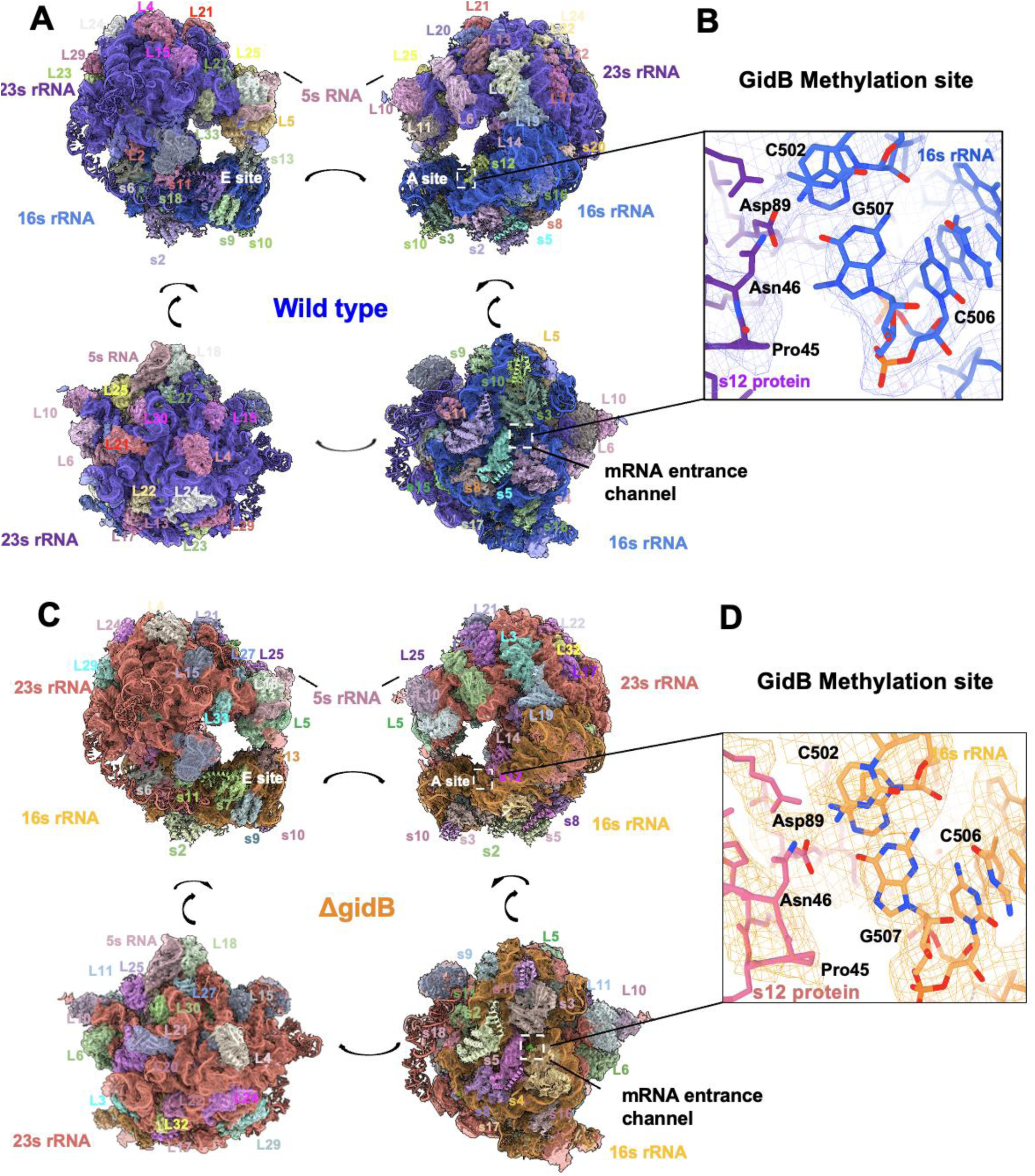
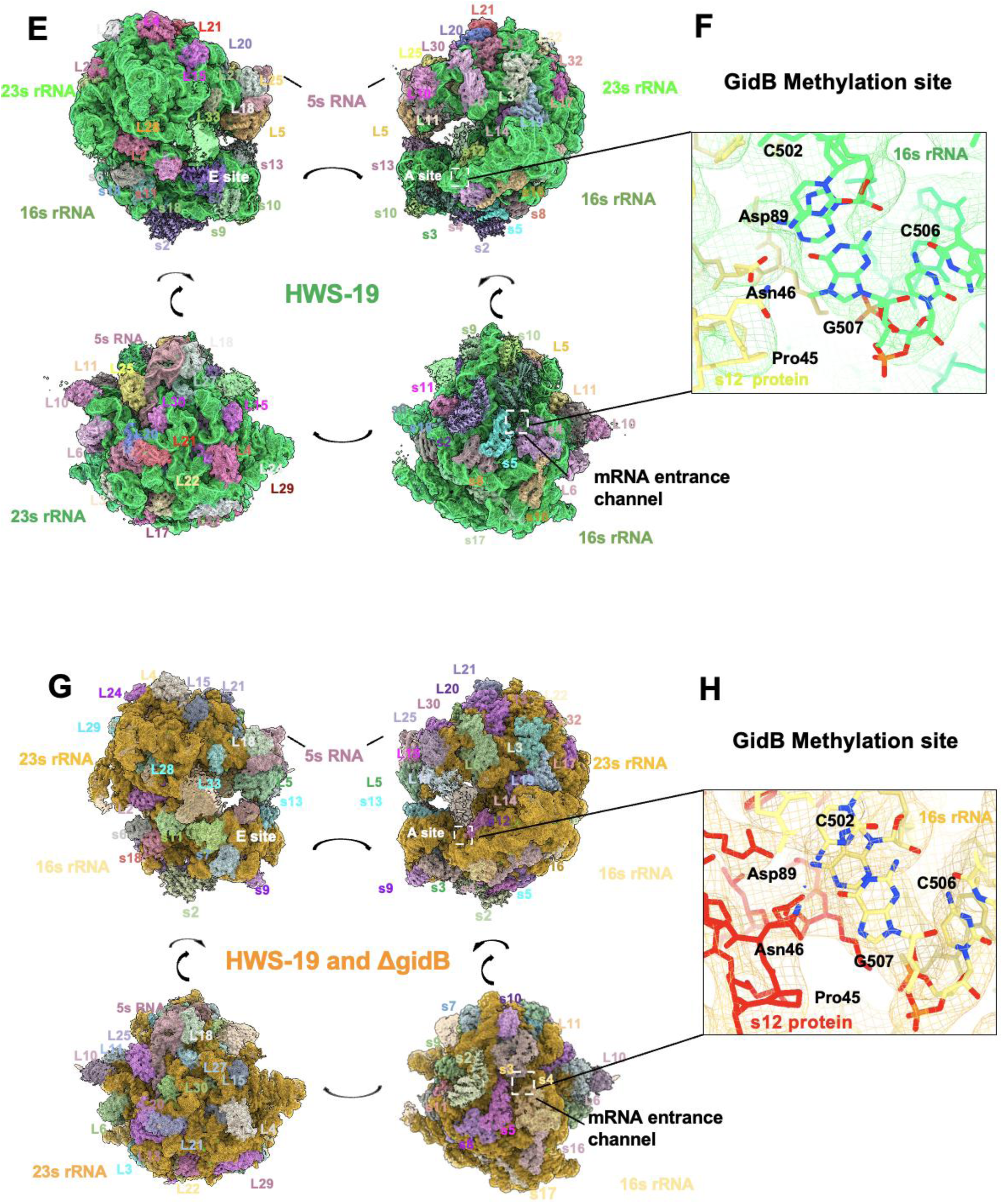
Structures of Mycobacterial Ribosomes with and without GidB-mediated methylation. Ribosomal structures from the following *Mycobacterium smegmatis* stains: wild type, ΔgidB, HWS-19 and HWS-19-ΔgidB strains. A) wild-type: Different views of the Msm-70S structure (particle number = 591,747, 3.47 Å resolution) and B) zoomed in site of the GidB methyltransferase target residue (G507). C) ΔgidB: Different views of the Msm-70S structure (particle number = 1,466,88, resolution = 2.47 Å) and D) zoomed in site of the GidB methyltransferase target residue (G507). E) HWS-19: Different views of the Msm-70S structure (particle number = 108,034, resolution = 3.26 Å) and F) zoomed in site of the GidB methyltransferase target residue (G507) and interacting residues. G) HWS-19-ΔgidB: Different views of the *Mycobacterium smegmatis* 70S ribosomal structure (particle number = 136,418, map resolution = 3.07 Å) and H) zoomed in site of the GidB methyltransferase target residue (G507) and interacting residues.

## Discussion

The ubiquitous presence of multiple and redundant proof-reading mechanisms for protein synthesis underscores the importance of accurate protein synthesis in cellular homeostasis. However, errors in protein synthesis are both more pervasive and frequent than previously anticipated, and increasingly, translational errors are being recognised as a mechanism for adaptation to hostile environments. One possible resolution of these two seemingly opposed processes is the recognition that ‘optimal’ fidelity is not to minimize errors to as low as possible and is context-specific [5, 6, 60]. For example, infection by *Salmonella* mutants with both impaired and enhanced translational fidelity were less productive than wild-type strains [26], and mycobacterial strains with extremely high mistranslation rates due to mutations in *gatA* grew more slowly than wild-type in axenic culture, but had orders of magnitude greater survival with rifampicin treatment [12].

In this study we sought to identify further translational fidelity factors in mycobacteria, which we had previously shown have high, but specific rates of mistranslation due to the indirect tRNA aminoacylation pathway required for aminoacylation of glutaminyl- and asparaginyl-tRNAs mediated by GatCAB [10, 12, 36]. Our screen identified loss-of-function mutations in *gidB* conferring increased fidelity to errors generated by mutants in this pathway (**Fig. 7**). We had previously shown that deletion of *gidB* in *M. tuberculosis* increased fidelity of ‘wobble’ ribosomal decoding errors, but not for mistranslation of asparagine for aspartate – one of the two specific errors in mycobacteria due to the indirect tRNA pathway [42]. Those studies, performed with otherwise wild-type mycobacteria under standard laboratory conditions, mimic similar conditions where in this study *gidB* deletion under those conditions also lacked a phenotype.

**Figure 7.**
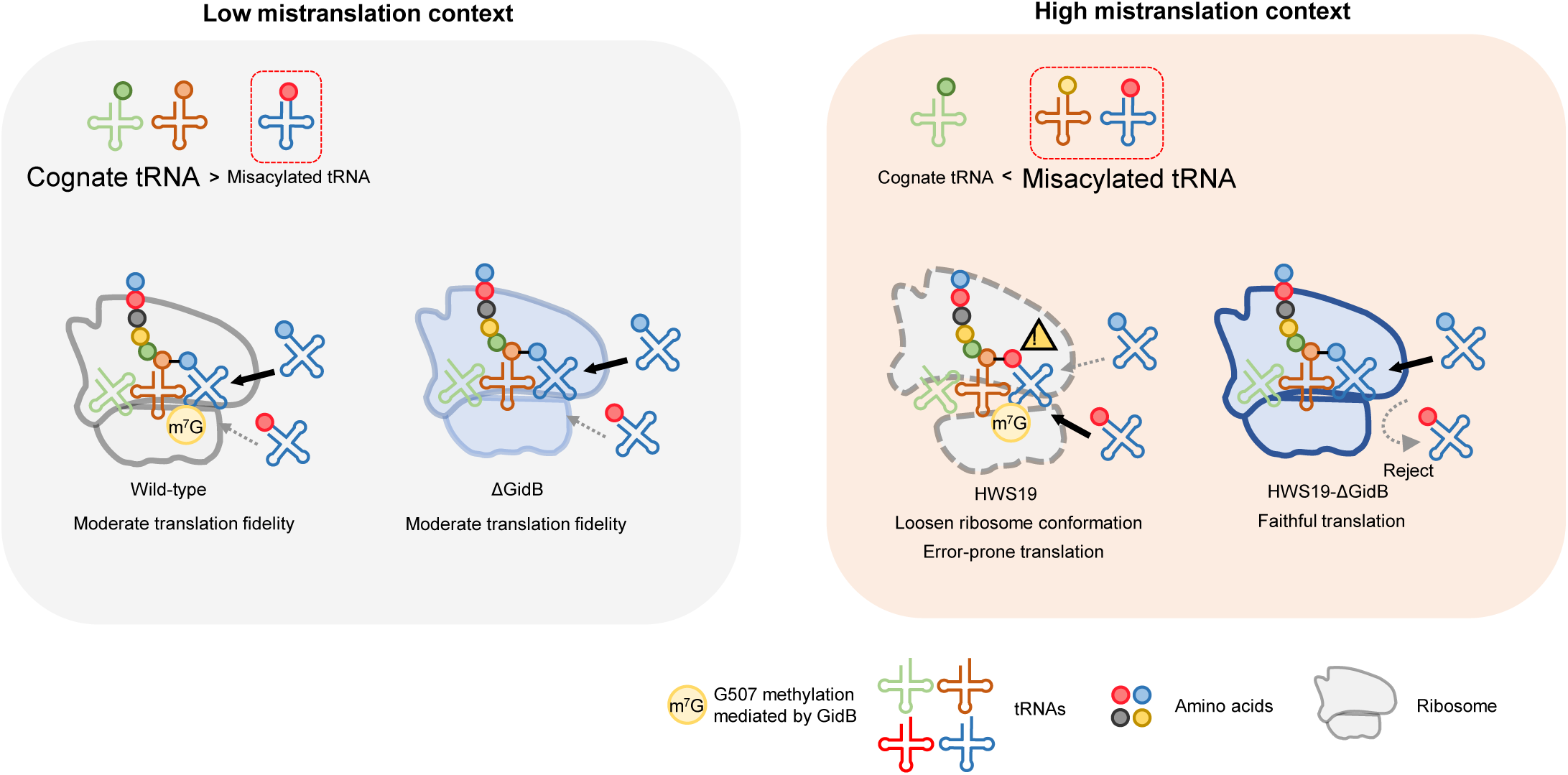
Proposed model of *gidB* mediated translation fidelity control. Under low mistranslation context (Left), wild-type ribosome and ΔGidB both have moderate translation fidelity. Under high mistranslation context (Right) due to impaired indirect aminoacylation pathway, HWS19 ribosome has a loosen ribosome conformation compared to HWS19-ΔGidB ribosome, which translation is error-prone. HWS19-ΔGidB ribosome restores structure integrity could better discriminate against misacylated-tRNA.

Although the ribosome is known to engage in a number of proof-reading functions, these have been confined to discrimination against non-cognate codon•anti-codon pairing, not discrimination of mischarged tRNAs where codon•anti-codon pairing remains fully cognate [2, 61–64]. One study by Cornish and colleagues specifically addressed the question of ribosome recognition of the aminoacyl identity of aa-tRNA by the *E. coli* ribosome [65]. Whilst they identified few differences, there was a subtle 2-3 fold increase in A-site sampling by mischarged tRNAs. *E. coli*, unlike mycobacteria and indeed most bacteria, does not routinely encounter mischarged tRNAs. Hence, this may suggest that other model systems may be better suited to investigate bacterial ribosome recognition and discrimination of mischarged tRNA. Our prior studies with the unusual aminoglycoside kasugamycin [10] also suggest ribosomes can discriminate against mischarged tRNAs. Kasugamycin was known to decrease ribosomal decoding errors [66, 67] in direct contrast with the increased errors witnessed with other aminoglycosides such as streptomycin. We showed that kasugamycin, at concentrations that did not inhibit protein synthesis, could decrease mistranslation from mischarged tRNAs not only in live mycobacteria, but also in a cell-free *in vitro* translation system, which had been modified to include excess mischarged Asp-tRNA^Asn^ [10].

We do not know the precise molecular mechanism by which deletion of *gidB* increases ribosomal discrimination of mischarged tRNAs. Our structural analysis suggests that there are no large changes in the overall conformational or composition of the ribosome upon deletion of *gidB.* However, it is important to remember that even though the HWS19-Δ*gidB* will lack the methylation at 16S G507, changes in translational fidelity relative to the HWS-19 ribosomes may result only from a small population of ribosomes that have certain asparagine-aspartate recoding events and even then, such effects may only be relevant at a specific step in the translation cycle. Our structural analysis identifies a likely candidate residue that could mediate this effect: S12 Asn46. Although both strains will have a small fraction of Asp46-containing S12 proteins, the increased discrimination of incorrectly charged tRNAs occurs only in the Δ*gidB* background. Previous work has shown how the deletion of *gidB* results in altered translational fidelity through interactions with the S12 loop that contains Asn/Asp46, which bridges 16S G507 to the codon:anticodon helix [42]. Collectively, these results suggest that a biochemically arrested complex containing a miscoded tRNA might result in structural differences in the HSW19-Δ*gidB* background only when S12 Asp46, but not Asn46, is present. These differences could reveal the mechanistic basis of the altered translational fidelity that was not resolvable here in empty ribosomes (without tRNA, mRNA, or nascent chain) containing a mixture of Asp/Asn at all positions. Future work may therefore require investigating several distinct Asp/Asn mutations and biochemical states to unravel whether changes to ribosomal composition or function are required for lack of GidB-mediated methylation to effectively discriminate against mischarged tRNAs.

We and others have recently described alternative bacterial ribosomes that are particularly adapted to stressful environments – such environments are also associated with relaxation of translational fidelity [32, 68, 69]. However, multiple forms of alternative ribosomes have been identified: comprising altered stoichiometry of ribosomal protein subunits, variant subunits or alternative rRNA composition [70, 71]. Which, if any of these alternatives contribute to increased fidelity in absence of GidB-mediated rRNA methylation will be the subject of ongoing study.

Deletion of *gidB* causes increased mycobacterial susceptibility to rifampicin. Since lack of *gidB* decreases both wobble misreading [42], as well as discrimination of mischarged tRNA, which function is responsible? Several lines of evidence suggest discrimination of mischarged tRNAs is more likely. First, we have previously demonstrated that increased discrimination of mischarged tRNA by kasugamycin is responsible for enhanced mycobacterial susceptibility to rifampicin *in vitro* [10]. Secondly, the magnitude of increased susceptibility of Δ*gidB* to rifampicin is greater in strain HWS19, in which the excess error is derived solely from mischarged tRNA. Finally, the absolute error rates of ribosomal decoding errors in mycobacteria, including wobble mistranslation, are much lower than for errors due to mischarged tRNA [17, 32, 33], making the latter mechanism more parsimonious.

Emerging data suggest absolute fidelity in protein synthesis is not desirable, not only from an efficiency perspective, but also because relaxed fidelity is associated with adaptation to hostile environments. Although moderate mistranslation rates may be adaptive under stress, excessive mistranslation is still harmful. Therefore, mechanisms that are permissive for moderate but not excessive mistranslation may serve an important function in bacterial environmental adaptation. Only mycobacteria with increased mistranslation rates – either via mutation or from an environmental context in which higher mistranslation rates are favoured – had a high fidelity phenotype in the absence of *gidB*. This extended to discrimination of mischarged tRNAs that are not naturally occurring, such as Ala-tRNA ^Ala^. Ribosomal RNA methylation may be reversible [72], although definitive evidence is not available. An analogy of the function/role of this system may be with an “automated braking system” in cars: i.e. it only comes into play at dangerous excessive speeds. Similarly, lack of methylation by GidB only prevents ‘runaway’ mistranslation that would otherwise result in error catastrophe, but allows for the potentially adaptive functions of translational error and is potentially tunable.

## MATERIALS AND METHODS

### Bacterial strains and culture

Wild-type *M. smegmatis* mc^2^-155 [73] and its derivatives were cultured in Middlebrook 7H9 Broth supplemented with 0.2% glycerol, 10% ADS (albumin-dextrose-salt) and 0.05% Tween-80 with corresponding antibiotics. Wild-type *E. coli* HK295 and its derivatives were cultured in LB Broth with corresponding antibiotics. *E. coli* DH5a and TOP10 were used for transformation of plasmids and amplification. LB-Agar was used for both *M. smegmatis* and *E. coli* in plating experiments. If not otherwise mentioned, bacteria were grown in 37 C, 220 rpm shaker or maintained in 37C incubator.

#### Suppressor screen

A total of around 1×10^7^ colony forming units (C.F.U) HWS19 were plated onto each of 6 LB agar plates containing 1μg/mL streptomycin. After 5 days, visible colonies were picked for further analysis. The *rpsL* gene was amplified by PCR and sequenced, those with wild-type *rpsL* were transformed with Renilla-Firefly dual luciferase reporter plasmids (Renilla-Firefly WT and Renilla-Firefly D214N). Specific mistranslation rates of asparagine for aspartate were measured in potential suppressor candidates to identify low mistranslating strains compared with HWS19. Genomic DNA was isolated from HWS19 and four suppressor candidates by standard methods, and were then whole-genomes sequenced and analyzed by Genewiz. The whole genome sequence data have been deposited to the NCBI short-read archive <https://www.ncbi.nlm.nih.gov/sra> with accession number: PRJNA704242. Mapping results of mutations onto genes covering over 99% of all reads in four suppressor candidates are shown as Table S1. A further 10 suppressor candidates had only *gidB* sequenced as per Table S2.

### Deletion of *gidB* and complementation in *Mycobacterium smegmatis*

The 500 bp upstream and downstream regions of *gidB* were amplified from mc^2^-155 genomic DNA. The zeocin resistance cassette was amplified from plasmid pKM-Zeo-Lox (A kind gift from Eric J. Rubin lab). Then the zeocin resistance cassette flanking 500 bp upstream and 500 bp downstream region of *gidB* were assembled by overlapping extension PCR for use as an allele exchange substrate (AES), which was verified by sequencing. WT mc^2^-155 and HWS19 transformed with pNIT(kan)::RecET::sacB recombineering plasmid were grown to OD_600_∼0.4, and expression of recET was induced with 10 μM isovaleronitrile (IVN) for 5 h, then competent cells were made by standard methods and transformed with 2 μg of the AES. The cells were recovered for 4 h and selected on LB agar plates containing 20 μg/mL zeocin. The recombinants were confirmed by PCR for verifying the zeocin cassette integration and the appropriate genomic context (Figure S3) (See verification primers sequences in Table S3). *gidB* deletion strains were then cured of the recombinase plasmid prior to further experiments.

For *gidB* complementation construction, several plasmids were constructed. The chromosome integrating mycobacterial plasmid pML1357 (Addgene #32378) was used as the backbone vector. The hygromycin resistance cassette was replaced with a kanamycin resistance cassette and streptomycin resistance cassette as pKML1357 and pSML1357 respectively. *gidB* from wild-type *M. smegmatis* with and without native promoter were amplified and cloned into integrative plasmids pKML1357/ pSML1357 to construct pKML1357-Pnative_*gidB*, pKML1357-Psmyc_*gidB*, pSML1357-Pnative-*gidB*. Then the plasmids were transformed into Δ *gidB* strains and selected onto LB agar with corresponding antibiotics for *gidB* complementation strains. Δ*gidB*::Pnative_*gidB* (KanR) strains were used for dual luciferase mistranslation assay when measuring N to D and Q to E mistranslation rates. Δ *gidB*::Psmyc_*MsmgidB* (KanR) were used for the dual-fluorescence mistranslation assay. Δ *gidB*::Pnative_*gidB* (StrepR) strains were used for dual-luciferase mistranslation assays when measuring W to A mistranslation as the pACET-Renilla-Firefly construct carries a kanamycin resistance marker. In addition, *gidB* mRNA levels in *gidB* deletion and complementation strains were verified by RT-qPCR (Figure S3).

### Measuring mistranslation rates with Renilla-Firefly dual-luciferase reporters

For measuring mistranslation rates in *M. smegmatis*, the shuttle plasmids pTetG-Renilla-Firefly (WT), pTetG-Renilla-D120N-Firefly, pTetG-Renilla-E144Q-Firefly under control of a tetracycline-inducible promoter were used as previously [17]. For measurement of tryptophan to alanine mistranslation rates, the mycobacterial chromosome integrating plasmid pACET-Renilla-Firefly, where expression of the reporter is under control of an acetamide-inducible promoter was used [17]. The *Renilla* gene was mutated by site-directed mutagenesis to Renilla-A214W.

In mycobacteria, the assay was performed as previously [12]. Briefly, strains with pTetG-Renilla-Firely were cultured to OD600>3, and diluted to OD600∼0.2 into fresh 7H9 medium supplemented with 50μg/ml hygromycin and 100 ng /ml anhydrotetracycline for inducing dual luciferase. After 7-8 hours, cells were collected by centrifugation at 12000rpm for 3min, lysed with 40μL passive lysis buffer in 96-well white flat bottom plate for 20-30min (plate shaking at 400rpm, room temperature), then reacted with 50μL substrate Firefly reagents, 360rpm shaked 5s followed by measuring the Firefly luminescence by Fluoroskan Ascent FL luminometer with 1000ms integration time. Then 50μL Stop&Glo reagent was added followed by measuring the Renilla luminescence immediately. For strains with pACET-Renilla-Firely and pTet-tRNA_CCA_^Ala^ [17], the strains in experimental group were diluted to OD600∼0.2 into fresh 7H9 medium supplemented with 50μg/ml hygromycin, 20% acetamide and 100ng/mL anhydrotetracycline for inducing dual luciferase and tRNA_CCA_^Ala^ respectively. The later procedures were same as above. The corrected Renilla/Firefly representing the mistranslation level is calculated as follows: Corrected Renilla / Firefly (N to D) = 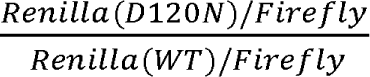, Corrected Renilla / Firefly (Q to E) = 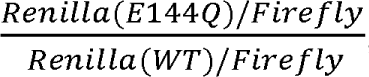, Corrected Renilla / Firefly (W to A)= 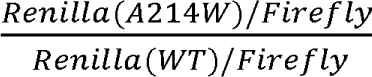.

### Measuring mistranslation rates with dual-fluorescence reporters

The pUVtetOR-GLR (green-linker-red) reporter was adapted as previously [12] with some modifications. Briefly, a mutated green fluorescent protein (E222Q) was fused to wild-type monomeric red fluorescent protein (mRFP) with a flexible GGSGGGGSGGGSSGG linker. A C-terminal tag (AANDENYAAAV) targeting the protein for proteolytic degradation [74] was added by PCR.

The procedure of measuring relative mistranslation rates by flow cytometry was as previously described [12] with some modifications. Strains with pUVtetOR-GLR were cultured to OD600∼0.2 and then 100ng/ml ATc was added for inducing dual-fluorescence. After 3 hours, cells were diluted into 1mL PBS to OD600∼0.05. The GFP and RFP signals of the sample were collected by flow cytometry (BD LSR Fortressa), being excited/ detected by 488nm/520nm laser line and 561nm/585nm laser line respectively. For the strains after high mistranslating selection: Strains expressing pSML1357-*Aph*_D214N and pUVtetOR-GLR were spread onto LB agar containing 50μg/ml Hygromycin and 2μg/ml Kanamycin as experimental group and LB agar containing only 50μg/ml Hygromycin as control group. After 5 days, colonies were scraped from the plates and cultured into 7H9 with 100ng/ml ATc for 3 hours. The following procedures were same as above. For the strains under rifampicin condition: strains with pUVtetOR-GLR were spread onto LB agar no selection plates and LB agar containing 20μg/ml rifampicin. After 5 days, colonies were then scraped from the plates and cultured into 7H9 with 100ng/ml ATc for 3 hours. The following procedures were same as above. Data were analyzed by FlowJo 10.4 for Windows 10. The gating strategy was as follows: Live bacteria were gated to eliminate debris as P1 based on SSC-A and FSC-A, then stringent gating on single cells was applied using a tandem gating strategy based on FSC-A and FSC-H as P2, subsequently using SSC-A and SSC-H as P3. Single cells with positive red fluorescence were acquired as P4 based on the red fluorescence signal. See Figure S4. Relative mistranslation rates were analysed as a histogram of GFP/ RFP ratio as P5 . The mean values of GFP/RFP ratio were calculated by FlowJo software (Flowjo 10.4 for Windows 10).

### Rifampicin time-kill curve assay

Wild-type and strain HWS19 *M. smegmatis* were cultured in 7H9 medium until OD600∼1. Cells were diluted to OD600∼0.3 into fresh 7H9 medium containing 20μg/ml rifampicin and cultured into 37°C, 220rpm shaker. Prior to addition of rifampicin, an aliquot was removed for calculation of cell numbers at time zero. At indicated time points, aliquots were taken, washed 3x in PBS and resuspended in PBS prior to being plated onto LB-agar. At least 3 10-fold dilutions were plated per time point. After 3-5 days, visible colonies were counted. The counts at different time points were normalized by counts at time zero for analysis.

### Relative Translation Rate Assay

To assess the relative translation rate, we adapted a previously described protocol [32] with minor modifications. *M. smegmatis* strains harboring the Nluc reporter construct was first grown in 7H9 medium to a stationary phase (ODLLL > 3). Cultures were then diluted to an ODLLL of 0.2 in fresh 7H9 medium, and anhydrotetracycline (ATc) was added to a final concentration of 50 ng/mL to induce reporter expression. After 5 minutes of induction, cells were pelleted, washed twice with fresh 7H9 medium (without ATc), and resuspended in the same medium supplemented with or without 200 μg/mL chloramphenicol (a translation inhibitor). Aliquots were collected every 10 minutes and immediately snap-frozen in liquid nitrogen. Once all time points were collected, luminescence was measured using a luminometer. All experiments were performed in triplicate using three independent biological replicates.

### 70S ribosome purification

HWS19 and HWS19-Δ*gidB* strains were grown in Middlebrook 7H9 Broth supplemented with 0.2% glycerol, 10% ADS (albumin-dextrose-salt) and 0.05% Tween-80 to OD 0.8. Bacteria pellet were collected by centrifuge at 4000rpm at 4C for 10 minutes. Pellet was washed and resuspend with polysome buffer (50 mM Tris-actate pH=7.2, 12 mM MgCl2, 50mMNH_4_Cl, 1mM DTT). Bacteria resuspension was lysed by MP FastPrep-24 bead beating system and the lysate was centrifuged at 12,000rpm 4C for 10 minutes. Cleared lysate was loaded on to 10% - 50% linear sucrose gradient with Beckman SW40Ti rotor at a speed of 39, 000rpm at 4C for 4 hours. Gradient were fractioned with continuous A260 measurement and the correspond 70S fraction were collected, buffer exchanged for downstream cryo-EM studies.

### CryoEM Data Collection and Map Refinement

Samples were diluted in milliQ ddH_2_O and deposited onto glow-discharged (EMS-100 Glow Discharge System, Electron Microscopy Sciences, 30 s at 15 mA and 22 mbar) copper Quantifoil (Quantifoil Micro Tools) grids, 1.2/1.3 spacing, 300-mesh, coated in 2 nm amorphous carbon. Grids were incubated for 30 s under 100% humidity and 10°C, blotted once at force 3 with Whatman #1 filter paper, and plunge-frozen in liquid ethane using a FEI Vitrobot Mark IV (Thermo Fisher). Grids were screened for ice quality on an FEI Talos Arctica (Thermo Fisher, 200 kV) or FEI Titan Krios (Thermo Fisher, 300 kV) transmission electron microscope at UCSF. WT strain non-mutant samples were shipped by dry shipper to the National Center for CryoEM Access and Training (NCCAT) at NYSBC. These were imaged on an FEI Titan Krios (Thermo Fisher, 300 kV) electron microscope with a Falcon IV camera using Leginon and without an energy filter. All other samples were imaged at UCSF on an FEI Titan Krios (Thermo Fisher, 300 kV) electron microscope with a Gatan K3 direct electron detector, using SerialEM and with an imaging filter (20 eV slit). Datasets at UCSF were collected with fringe-free imaging using a nine-shot beam-image shift approach with coma compensation. Further details are reported in Table 1. For all datasets, image stacks were binned by a factor of 2, motion corrected, and dose weighted using UCSF MotionCor2. Contrast transfer function (CTF) initial parameters were determined using CTFFIND4 in cisTEM (development version). After excluding images with poor CTF fits or poor ice quality, dose-weighted micrographs were subjected to unsupervised particle picking with a soft-edged disk template. Two-dimensional classification of the resulting particle stacks was used to select the images carried forward. Ab initio reconstructions were carried out for each dataset to produce initial references for three-dimensional reconstruction, which proceeded through auto refinement and manual refinement in two to three classes as necessary to exclude non-ribosome particles and 50S subunits. Final reconstructions were generated without sharpening from the classes yielding the 70S ribosome in the latest (highest resolution) refinement cycle prior to the emergence of overfitting artifacts in the Fourier shell correlation (FSC). Map resolutions are reported as particle FSC at 0.143.

**Table 1.** Cryo-EM data collection, refinement and validation statistics.

|  | WT strain WT<br>(EMDB-44092)<br>(PDB 9B1Y) | WT strain gidB<br>mutant<br>(EMDB-44097)<br>(PDB 9B24) | HWS19 strain<br>WT<br>(EMDB-44090)<br>(PDB 9B1W) | HWS19 strain<br>gidB mutant<br>(EMDB-44091)<br>(PDB 9B1X) |
| --- | --- | --- | --- | --- |
| **Data collection and processing** |  |  |  |  |
| Magnification | 96000 | 105000 | 130000 | 130000 |
| Voltage (kV) | 300 | 300 | 300 | 300 |
| Electron exposure (e-/Å<sup>2</sup>) | 60.74 | 901.165 | 72.704 | 72.704 |
| Defocus range (μm) | 400-1400 | 500-1500 | 500-1500 | 500-1500 |
| Pixel size (Å) | 0.833 | 0.833 | 0.662 | 0.664 |
| Symmetry imposed | P1 | P1 | P1 | P1 |
| Initial particle images (no.) | 704502 | 1636851 | 254392 | 417089 |
| Final particle images (no.) | 591747 | 1466881 | 108034 | 136418 |
| Map resolution (Å) | 3.47 | 2.47 | 3.26 | 3.07 |
| FSC threshold | 0.143 | 0.143 | 0.143 | 0.143 |
| Map resolution range (Å) | inf-3.47 | inf-2.47 | inf-3.26 | inf-3.07 |
| **Refinement** |  |  |  |  |
| Initial model used (PDB code) | 5o60, 505j | 5o60, 505j | 5o60, 505j | 5o60, 505j |
| Model resolution (Å) | 3.32 | 3.05 | 4.22 | 3.69 |
| FSC threshold | 0.143 | 0.143 | 0.143 | 0.143 |
| Model resolution range (Å) | inf-3.47 | inf-2.47 | inf-3.26 | inf-3.07 |
| Map sharpening *B* factor (Å<sup>2</sup>) | N/A | N/A | N/A | N/A |
| **Model composition** |  |  |  |  |
| Non-hydrogen atoms | 146981 | 146980 | 144760 | 144818 |
| Protein residues | 5723 | 5723 | 5877 | 5877 |
| Ligands | 629 | 629 | 629 | 629 |
| ***B* factors (Å<sup>2</sup>)** |  |  |  |  |
| Protein | 140.30 | 197.35 | 139.43 | 191.74 |
| Ligand | 142.41 | 162.55 | 171.92 | 154.91 |
| **R.m.s. deviations** |  |  |  |  |
| Bond lengths (Å) | 0.014 | 0.004 | 0.003 | 0.010 |
| Bond angles (°) | 0.893 | 0.670 | 0.541 | 0.650 |
| **Validation** |  |  |  |  |
| MolProbity score | 2.72 | 2.08 | 1.80 | 1.70 |
| Clashscore | 59.82 | 19.66 | 9.34 | 7.56 |
| Poor rotamers (%) | 0.86 | 0.13 | 0.40 | 0.10 |
| **Ramachandran plot** |  |  |  |  |
| Favored (%) | 92.44 | 95.78 | 95.56 | 95.82 |
| Allowed (%) | 7.47 | 4.17 | 4.36 | 4.11 |
| Disallowed (%) | 0.09 | 0.05 | 0.09 | 0.07 |

### Atomic Structure Refinement

An initial atomistic model of the 70S ribosome was generated by fitting the 5o5j and 5o60 PDB models of the small and large ribosomal subunits in Mycobacterium smegmatis, respectively, into the final map of the HWS19 unmutated dataset using UCSF ChimeraX. This model was then subjected to several iterations of rigid body fitting in UCSF ChimeraX or PyMOL, molecular dynamics model fitting in ISOLDE, model building/trimming/correction in Coot or UCSF ChimeraX, and real-space model refinement with anisotropic displacement parameter (ADP) refinement in Phenix, not necessarily in that order, until convergence. Restraints for residues not distributed with Phenix libraries were generated using Schrödinger and Phenix. The final HWS19 unmutated model was re-fitted and re-refined into the other three maps, with further model trimming as needed where map quality was not sufficient to resolve the structure. Figures were prepared using the PyMOL Molecular Graphics System (Schrödinger).

### Statistical analysis

All statistical methods are described in figures legends, presented as mean with SD. The p values and statistical significance were calculated using GraphPad prism software (Prism 8 for Mac). Two-tailed unpaired Student’s t test was used to compare means between groups. ***p<0.001, **p<0.01, *p<0.05, ns p>0.05.

### Author contributions

HWS optimised the conditions of and performed the initial screen. ZB performed most experiments with assistance from JYH and YC. YXC purified the 70S ribosomes, interpreted data and modified all figures. HJ performed additional experiments for the revised manuscript. ZB and YXC analysed the results. IYand JSF solved the cryo-EM structure and MTD and JSF revised the analysis and prepared revised structural figures. BJ conceived of and supervised the study. ZB, YXC and BJ wrote the manuscript with input from the other authors.

## Acknowledgements

We would like to thank the Tsinghua flow cytometry core for technical assistance. We would like to thank Dr. Peter Walter at UCSF use of the ultracentrifuge and the fractionator in his lab. We appreciate the members from the Javid Lab and Fraser Lab for discussion and feedback on this study. This study was in part funded by grants from the Bill & Melinda Gates Foundation (OPP1109789) and funds from Tsinghua University School of Medicine to BJ and a Wellcome Trust Investigator award (207487/C/17/Z) and NIAID R56AI185094 to BJ (for work performed at UCSF), and NIH R35GM145238 to JSF. The Cryo-EM equipment at UCSF is partially supported by NIH grants S10OD020054, S10OD021741 and S10OD026881.

