## Supplementary material for "Ribosomal RNA methylation by GidB modulates discrimination of mischarged tRNA": Supp information

### Supplementary information-Figures

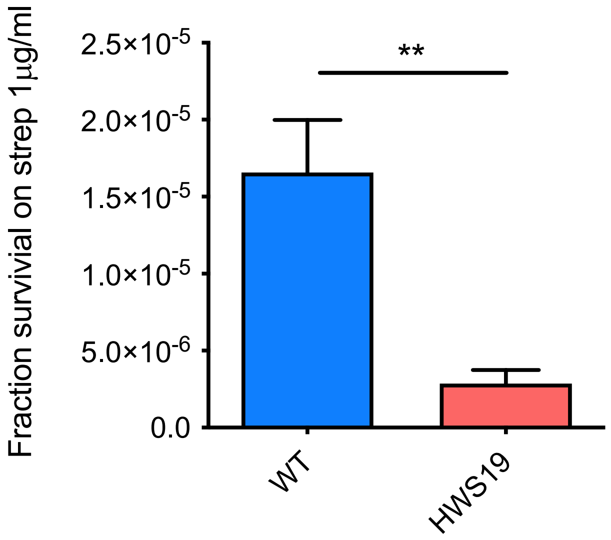


**Figure S1. Susceptibility to streptomycin of WT and HWS19**. HWS19 is more susceptible to MIC of streptomycin than WT. Data are presented as means of three biological replicates ± SD. **P<0.01

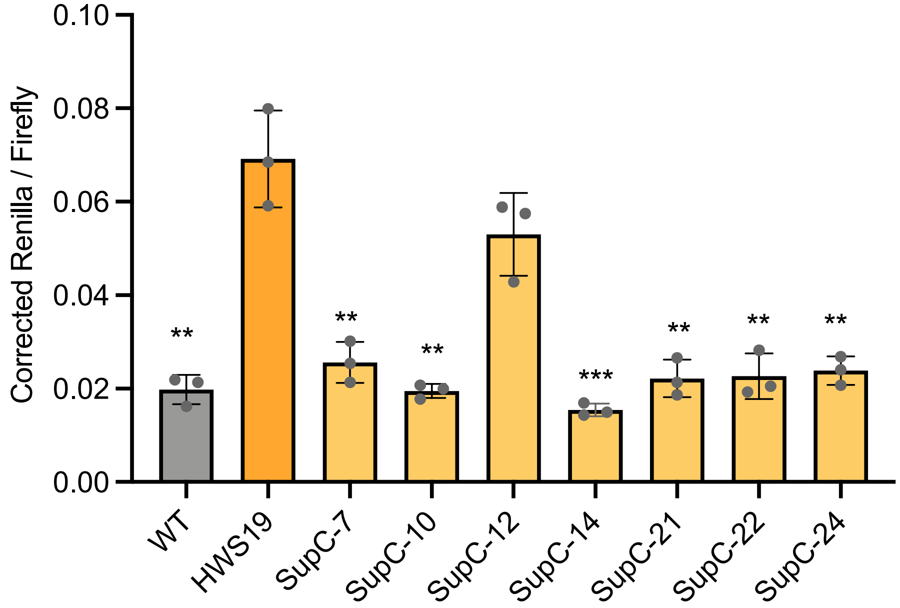


**Figure S2. Mistranslation level of the other suppressor candidates with *gidB* mutations.** N to D mistranslation rates were measured in 7 suppressor candidates compared with wild type and HWS19 by Renilla-Firefly dual luciferase reporter. Mutations in each suppressor candidates were shown in Table S2.

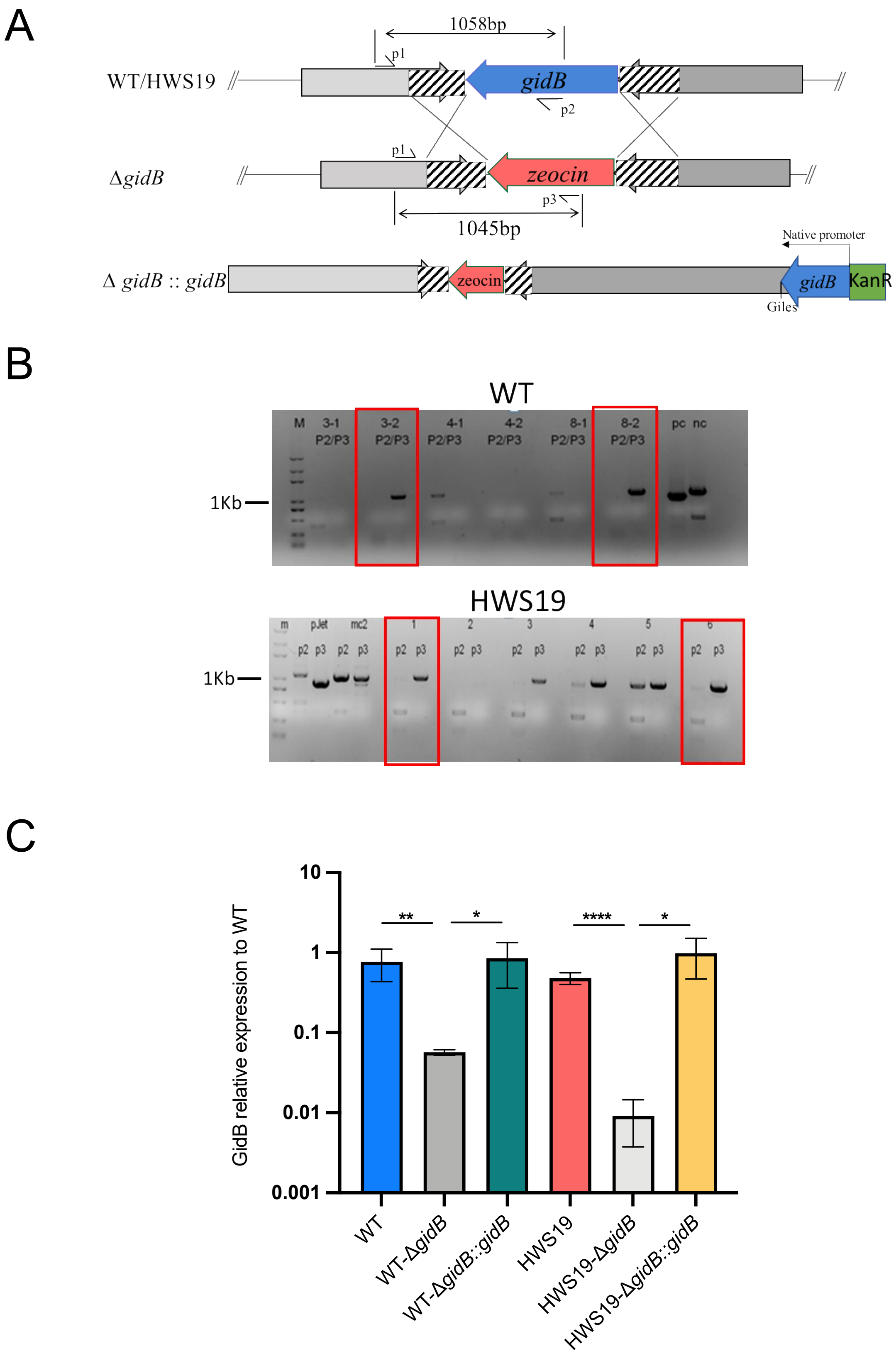


**Figure S3. *gidB* deletion and complementation in WT and HWS.19.** A.Homologous recombination was performed for deletion of *gidB,* replaced with zeocin resistance marker*. gidB* complement was integrated into Giles site with pml1357. B. PCR screening to verify successfully deletion of *gidB* in both wildtype background and HWS19 backgroud. Primer 1 (P1) is upstream of *gidB*, primer 2 (P2) is inside of *gidB*, primer 3 (P3) in inside of zeocin resistance marker. See Table S3 for sequences of primers. Red box means successfully deletion of *gidB*. C. RT-qPCR to further verify the *gidB* mRNA level in *gidB* deletion strains and complementation strains.

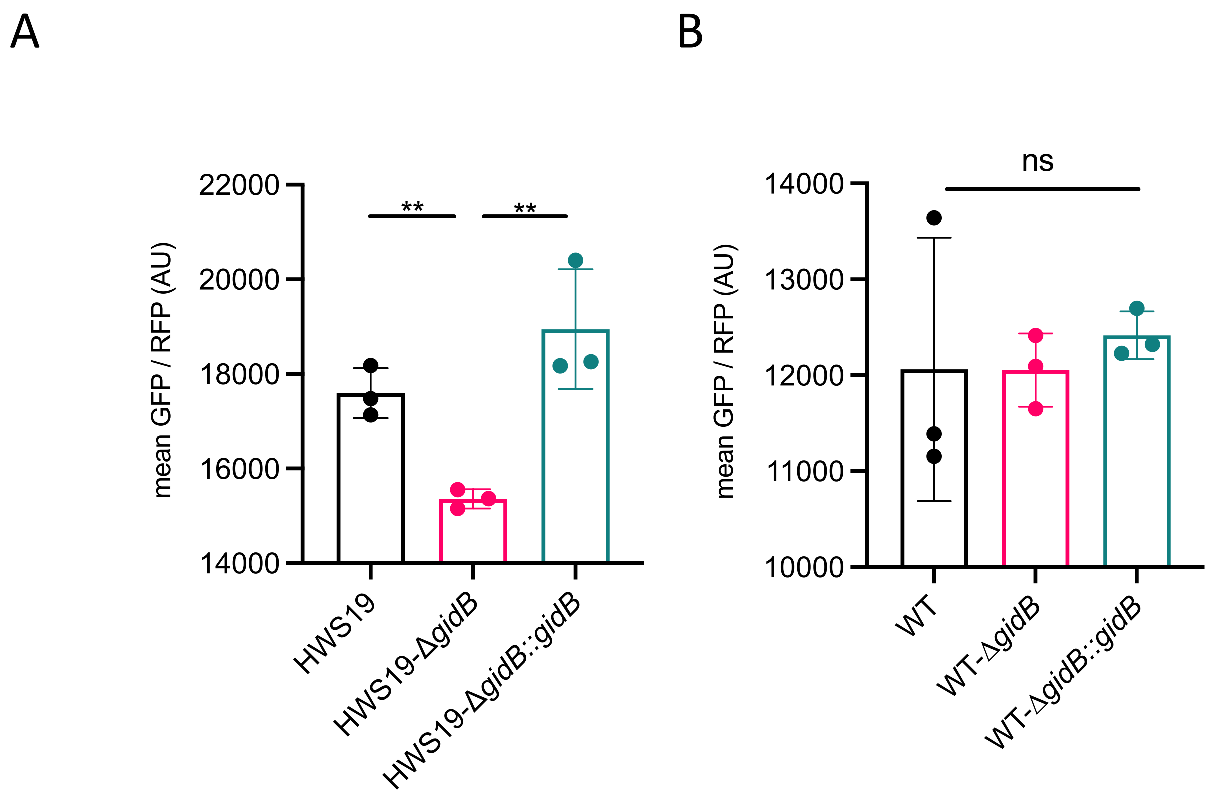


**Figure S4. *gidB* deletion increases translation fidelity in high mistranslation mutant.**
E to Q mistranslation measured by gain-of-function dual fluorescent reporter (Figure.2B) in high mistranslation HWS19 strain (A) and wild type strain (B)

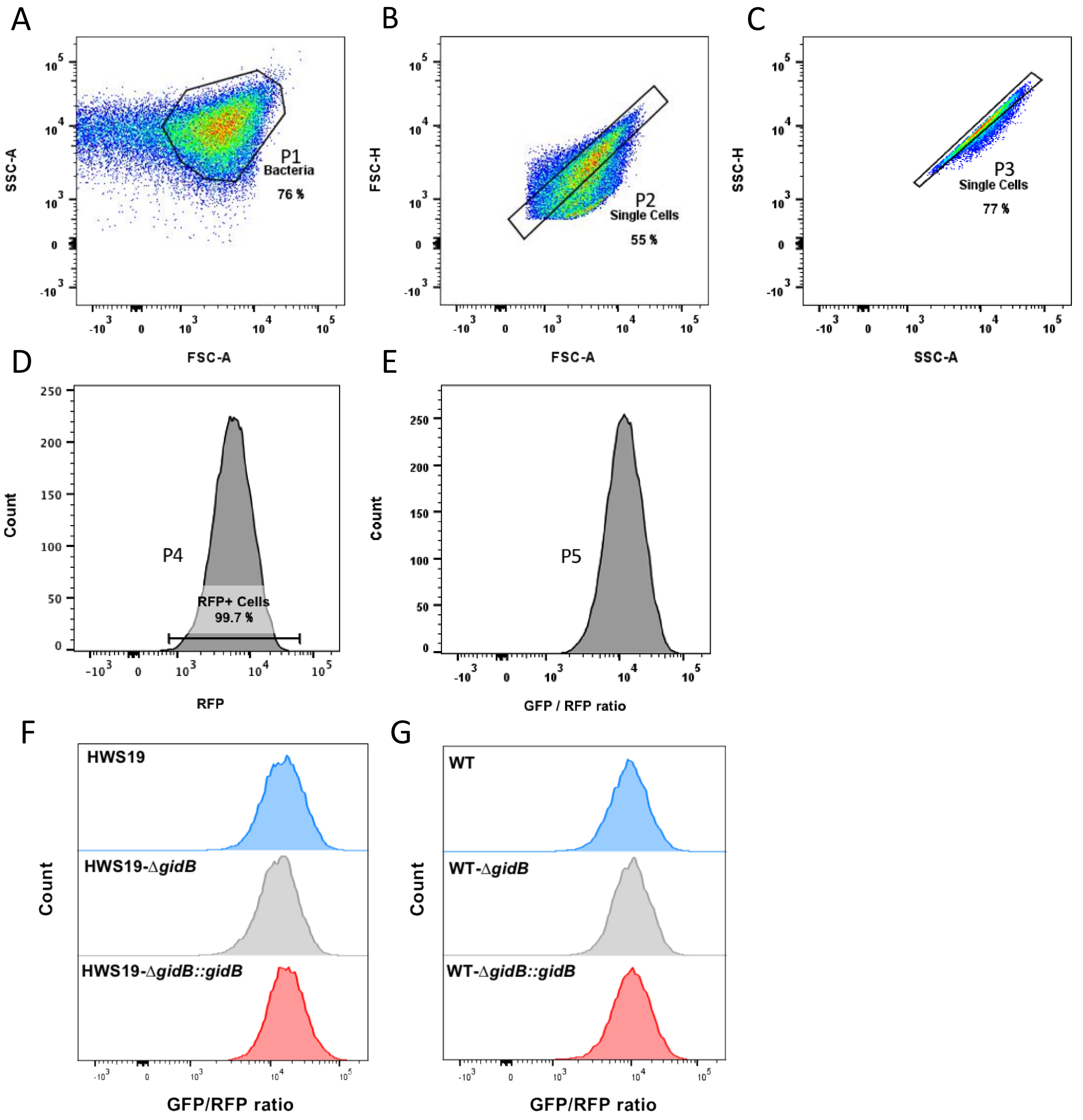


**Figure S5. Gating strategy of flow cytometry and histogram of population count replying on GFP/RFP ratio. A**. Gating *M.smegmatis* based on the estimation of the size and granularity as P1 in terms of SSC-A (X axis) and FSC-A (Y axis). B. Gating single-cell as P2 in terms of FSC-A (X axis) and FSC-H (Y axis). C. Gating single-cell again as P3 in terms of SSC-A (X axis) and SSC-H (Y axis). D. Gating smegmatis expressing positive RFP as P4 in terms of RFP (X axis). E. Histogram of population count (Y axis) relying on GFP/RFP ratio (X axis). F.G. GidB deletion strains and complementation strains in high mistranslating background (F) and wild-type background (G). X axis is the value of GFP and RFP ratio, representing the Q to E mistranslation level. The strain distributes to the right presents a higher mistranslation level than the one to the left.

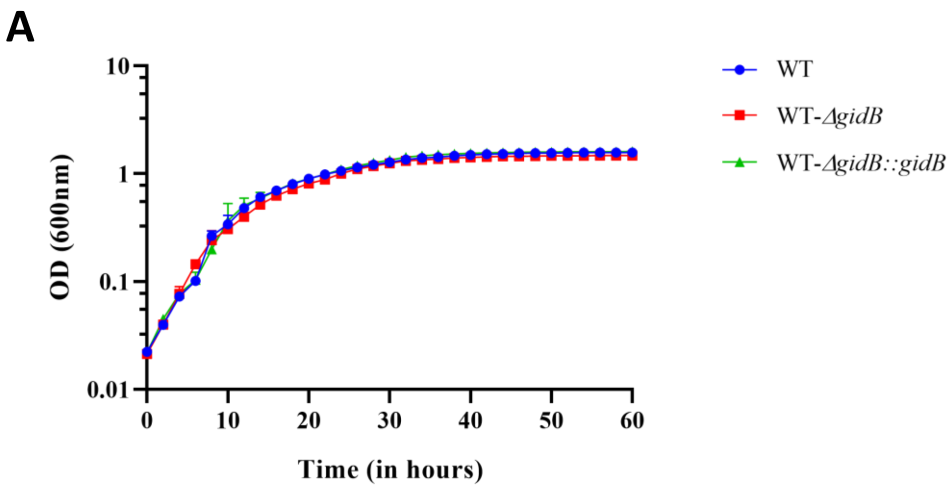


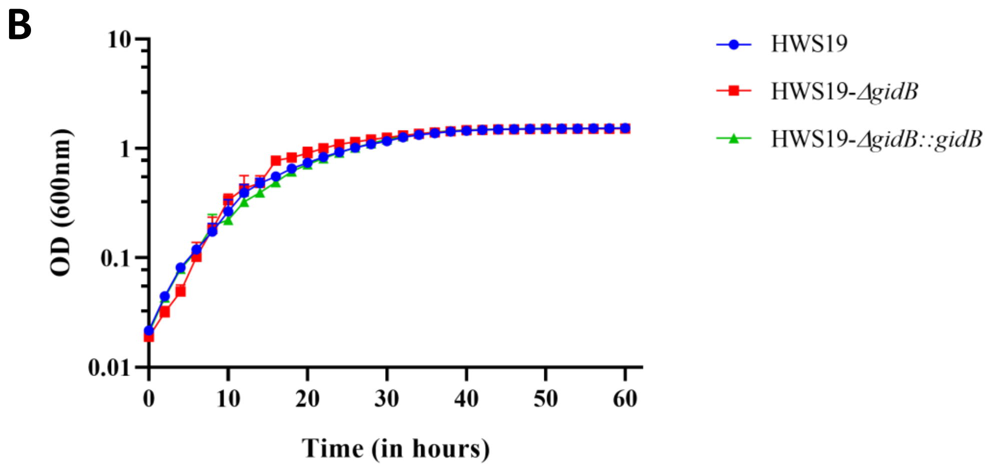


**Figure S6. Deletion of GidB does not affect growth rate in axenic culture.** WT Msm and Δ*gidB* (A) and strain HWS19 and HWS19Δ*gidB* (B) and their respective complemented strains were grown in standard supplemented 7H9 medium and optical density monitored over time. Points represent means +/- SD.

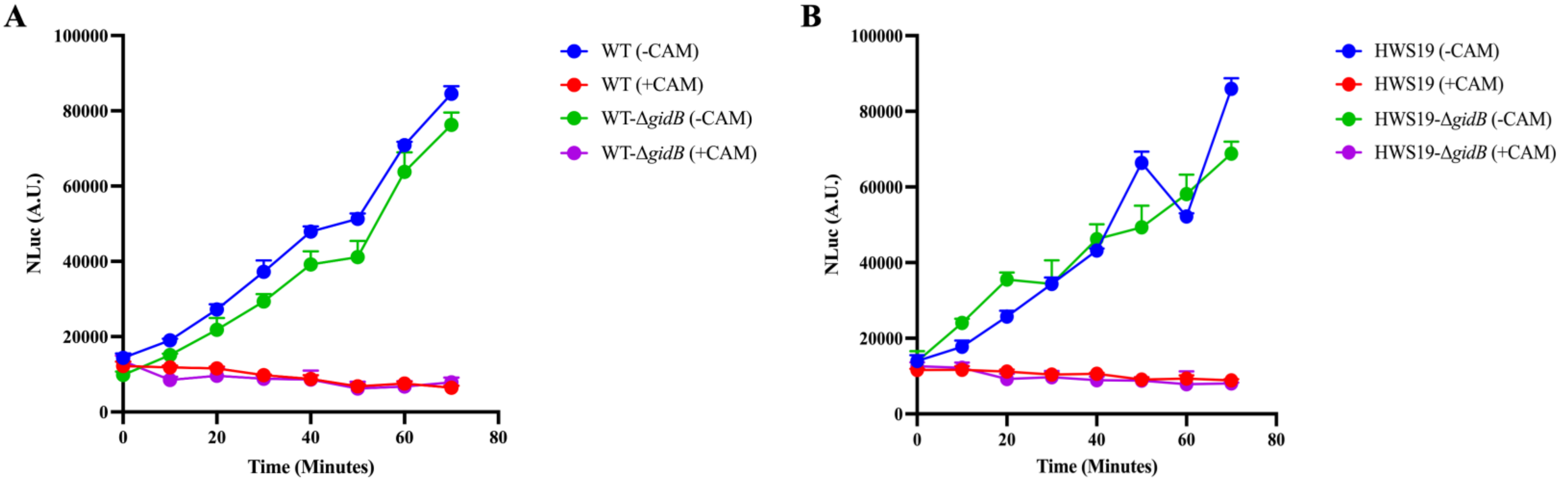


**Figure S7. Deletion of GidB does not affect translation rate of Nluc luciferase.** WT Msm and Δ*gidB* (A) and strain HWS19 and HWS19Δ*gidB* (B) were transformed with a tetrycline-inducible Nluc luciferase construct. Nluc activity was monitored over time as a proxy for translation rate in all 4 strains. Time =0 represents time of addition of Atc (50ng/mL). Experiment was performed using three independent biological replicates. Points represent means +/- SD. Chloramphenicol (200 µg/mL) was used as a translation inhibitor control.

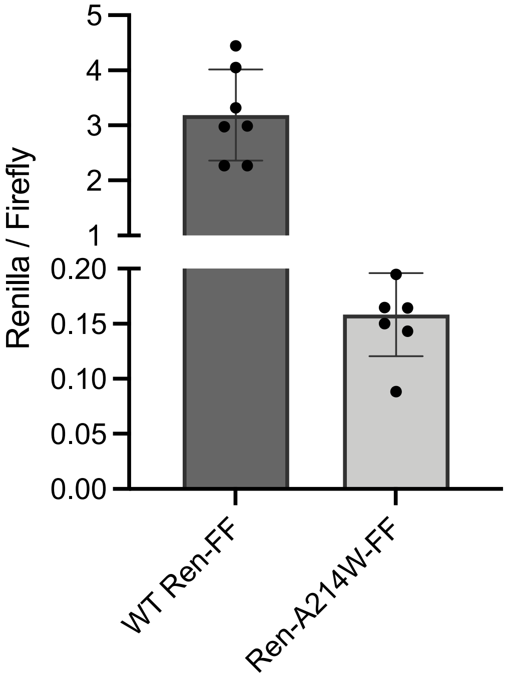


**Figure S8. Corrected Renilla luciferase activity.** Renilla-WT_Firefly and Renilla-A214W_Firefly were transformed into wildtype smegmatis. The absolute ratio of Renilla-WT/ Firefly and Renilla-A214W/ Firefly were illustrated in the graph.

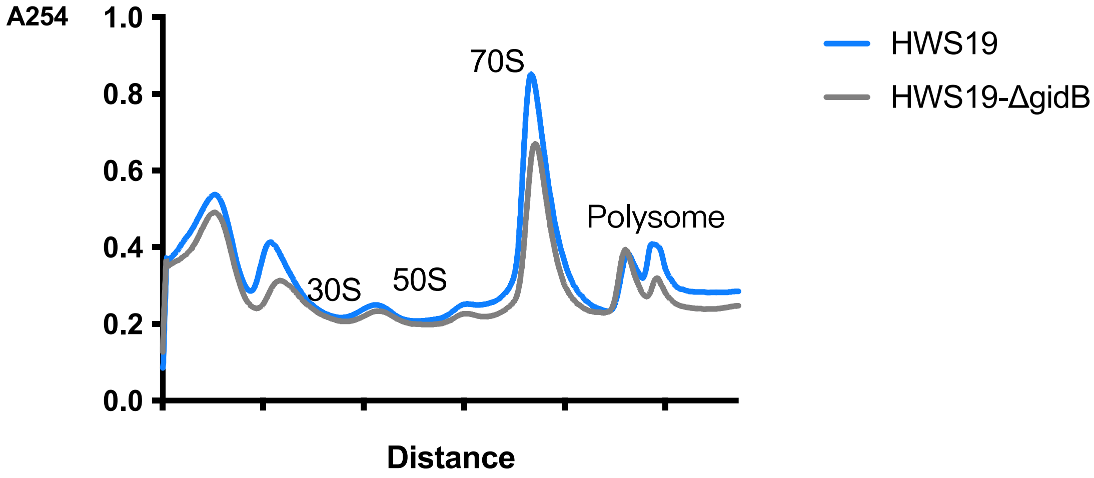


**Figure S9. Subunit profiling of different *M.smegmatis* strains.** Sucrose gradient profile of HWS19, HWS19- ΔgidB.
